# Drought stress delays photosynthetic induction and accelerates photoinhibition of photosystem I under fluctuating light

**DOI:** 10.1101/2021.10.07.463522

**Authors:** Hu Sun, Qi Shi, Ning-Yu Liu, Shi-Bao Zhang, Wei Huang

**Author notes:** **Corresponding authors:** Shi-Bao Zhang, Wei Huang. **Funding:** 1. the National Natural Science Foundation of China (No. 31971412, 32171505). 2. the Project for Innovation Team of Yunnan Province.

## Abstract

Fluctuating light (FL) and drought stress usually occur concomitantly. However, whether drought stress affects photosynthetic performance under FL remains unknown. Here, we measured gas exchange, chlorophyll fluorescence, and P700 redox state under FL in drought-stressed tomato (*Solanum lycopersicum*) seedlings. Drought stress significantly affected stomatal opening and mesophyll conductance after transition from low to high light and thus delayed photosynthetic induction under FL. Therefore, drought stress exacerbated the loss of carbon gain under FL. Furthermore, restriction of CO_2_ fixation under drought stress aggravated the over-reduction of photosystem I (PSI) upon transition from low to high light. The resulting stronger FL-induced PSI photoinhibition significantly supressed linear electron flow and PSI photoprotection. These results indicated that drought stress not only affected gas exchange under FL but also accelerated FL-induced photoinhibition of PSI. Furthermore, drought stress enhanced relative cyclic electron flow in FL, which partially compensated for restricted CO_2_ fixation and thus favored PSI photoprotection under FL. Therefore, drought stress has large effects on photosynthetic dark and light reactions under FL.

## Introduction

During diurnal photosynthesis under natural field conditions, leaves usually experience dynamic changes in light intensity on timescales from milliseconds to hours, owing to the variation of the solar angle, cloud movement, wind-induced leaf fluttering, and shading from overlapping leaves and neighboring plants (Pearcy 1990; Slattery, Walker, Weber & Ort 2018). In addition to fluctuating light (FL), plants often experience suboptimal conditions, such as drought. Many studies have examined the effect of drought stress on photosynthesis under stable light intensity (Oukarroum, Schansker & Strasser 2009; Galmés *et al*. 2013; Zivcak *et al*. 2013). However, drought stress is rarely studied under FL conditions (Grieco *et al*. 2020; Qu *et al*. 2020). Under FL, plants may exhibit different photosynthetic performance when also experiencing drought stress compared with sufficient water conditions. To understand how plants perform under a combination of FL and drought stress in the field, it is important to explore the photosynthetic physiology of drought-stressed plants under FL.

After transition from low to high light, the net CO_2_ assimilation rate (*A*_N_) gradually increases, which is termed “photosynthetic induction”. The rate of photosynthetic induction significantly affects carbon gain and thus affects plant biomass when grown under FL (Adachi *et al*. 2019; Kimura, Hashimoto-Sugimoto, Iba, Terashima & Yamori 2020; Yamori, Kusumi, Iba & Terashima 2020). Therefore, promoting the rate of photosynthetic induction is a potential target for improving crop yield (De Souza, Wang, Orr, Carmo-Silva & Long 2020; Acevedo - Siaca *et al*. 2020). Under high light, *A*_N_ is determined by CO_2_ diffusional conductance and biochemical factors (Grassi & Magnani 2005). CO_2_ diffusional conductance refers to CO_2_ diffusion from the atmosphere to the chloroplast stroma, including stomatal conductance (*g*_s_) and mesophyll conductance (*g*_m_) (Flexas *et al*. 2014). Biochemical factors include the activation of electron transport, Calvin–Benson cycle enzymes, especially Rubisco, and sucrose synthesis (Sakoda, Yamori, Groszmann & Evans 2021). A recent study revealed that the induction of *A*_N_ under FL was mainly affected by *g*_s_ rather than *g*_m_ in *Arabidopsis thaliana* and tobacco (*Nicotiana tabacum*) plants in the absence of water stress (Sakoda *et al*. 2021). Increasing the stomatal opening significantly enhanced the induction of *A*_N_ and plant biomass under FL (Kimura *et al*. 2020; Yamori *et al*. 2020). Drought stress usually reduces *g*_s_ and *g*_m_ under constant high light, leading to decreased chloroplast CO_2_ concentration (*C*_c_) (Warren, Livingston & Turpin 2004). Under such conditions, CO_2_ fixation in chloroplast carboxylation sites is restricted by the lack of CO_2_. Therefore, the relative limitations of *g*_s_ and *g*_m_ imposed on *A*_N_ are enhanced under drought stress. However, it is unclear whether drought stress affects the induction responses of *g*_s_, *g*_m_, and *A*_N_ under FL. Characterizing the dynamics of *g*_s_ and *g*_m_ and how they limit *A*_N_ under drought and FL is crucial for understanding the physiological mechanisms regulating carbon gain by plants under suboptimal field conditions.

In addition to photosynthetic dark reactions, FL affects photosynthetic light reactions. Upon a sudden increase in light intensity, electron flow from photosystem II (PSII) to photosystem I (PSI) rapidly increases (Huang, Yang & Zhang 2019a; Sun, Zhang, Liu & Huang 2020b). Meanwhile, CO_2_ fixation is not fully activated (Tanaka, Adachi & Yamori 2019), generating an imbalance between the production of excited states and the consumption of reducing power (Gerotto *et al*. 2016; Tan, Huang, Zhang, Zhang & Huang 2021). The resulting PSI over-reduction increases the production of reactive oxygen species (ROS) in PSI. Consequently, oxidative damage to PSI occurs because the ROS produced within thylakoid membranes cannot be immediately scavenged by antioxidant systems (Takagi, Takumi, Hashiguchi, Sejima & Miyake 2016). Therefore, FL can cause selective photoinhibition of PSI in many angiosperms (Yamori, Makino & Shikanai 2016; Yamamoto & Shikanai 2019; Huang, Yang & Zhang 2019b). PSI damage suppresses photosynthetic electron flow, photoprotection, and CO_2_ assimilation (Sejima, Takagi, Fukayama, Makino & Miyake 2014; Zivcak, Brestic, Kunderlikova, Sytar & Allakhverdiev 2015; Shimakawa & Miyake 2019), first impairing plant growth and even causing plant death (Suorsa *et al*. 2012; Yamori *et al*. 2016). The PSI redox state under FL is significantly affected by the electron sink downstream of PSI (Wada *et al*. 2018; Tazoe *et al*. 2020; Sun, Yang & Huang 2020a). However, little is known about the effects of drought stress on the PSI redox state and PSI photoinhibition under FL.

Plants have evolved several alternative electron flows to protect PSI against photoinhibition under FL (Allahverdiyeva, Suorsa, Tikkanen & Aro 2015; Armbruster, Correa Galvis, Kunz & Strand 2017; Shikanai & Yamamoto 2017). Cyclic electron flow (CEF) around PSI is used by angiosperms to fine-turn photosynthetic apparatus under FL (Suorsa *et al*. 2012; Yamamoto & Shikanai 2019). After transition from low to high light, CEF activity usually increases to allow fast formation of the trans-thylakoid proton gradient (ΔpH), which alleviates the PSI over-reduction at donor and acceptor sides (Kono, Noguchi & Terashima 2014; Yang, Ding & Huang 2019a). Once a sufficient ΔpH forms, CEF activity decreases to steady-state levels to prevent over-acidification of the thylakoid lumen. Therefore, the dynamic regulation of CEF activity under FL is crucial for balancing photoprotection and photosynthesis (Alboresi, Storti & Morosinotto 2019). Previous studies have documented that CEF activity under FL is largely affected by the redox state of PSI (Yang *et al*. 2019a; Tan *et al*. 2021). In particular, CEF activity increases with moderate PSI over-reduction but maintains at a low level when PSI over-reduction is missing. However, the effects of drought stress on the dynamic regulation of CEF activity under FL is largely unknown.

The aim of this study was to investigate whether and how drought stress affects photosynthetic light and dark reactions under FL. With this knowledge it may be possible to understand how drought stress interacts with FL to affect photosynthetic physiology and plant growth. Tomato was used in this study, as it is a C3 model species with intermediate leaf photosynthetic capacity and an important vegetable crop worldwide. To address the above question, tomato plants were grown under full sun light with sufficient or deficient water. We then measured the rapid changes in gas exchange, chlorophyll fluorescence, and P700 signals under FL.

## Materials and methods

### Plant materials and growth conditions

Tomato (*Solanum lycopersicum* Miller cv. Hupishizi) plants were grown in a greenhouse under 40% full sunlight. Day/night air temperatures were approximately 30/20ºC, relative humidity was approximately 60-70%, and maximum light intensity was approximately 800 μmol photons m^−2^ s^−1^. The plants were grown in 19-cm diameter plastic pots with humus soil, and the initial soil N content was 2.1 mg/g. Plants were fertilized with 0.15 g N/plant every two days using Peters Professional’s water solution (N:P:K = 15:4.8:24.1) and watered every day. After cultivation for one and a half months, plants were watered using running water with 400g/pot (CK) or 200g/pot (drought) for 1 week, and then mature leaves were used for photosynthetic measurements.

### Chlorophyll content measurement in vivo

A handheld chlorophyll meter (SPAD-502 Plus; Minolta, Tokyo, Japan) was used to non-destructively measure the relative content of chlorophyll per unit leaf area.

### Gas exchange and chlorophyll fluorescence measurements

An open gas exchange system (LI-6400XT; Li-Cor Biosciences, Lincoln, NE, USA) was used to simultaneously measure gas exchange and chlorophyll fluorescence. After photosynthetic induction at 1500 μmol photons m^−2^ s^−1^ and 400 μmol mol^−1^ CO_2_ concentration for 20 min, the net CO_2_ assimilation rate and *g*_s_ reached steady-state values. Then, the light intensity was changed to 100 μmol photons m^−2^ s^−1^ for 5 min to conduct low light adaptation. Afterward, the light intensity was changed back to 1500 μmol photons m^−2^ s^−1^ to measure the photosynthetic induction phase. After adequate photosynthetic induction, the response of CO_2_ assimilation rate to incident intercellular CO_2_ concentration (*A*/*C*_i_) curves were measured by decreasing the CO_2_ concentration to a lower limit of 50 μmol mol^−1^ and then increasing stepwise to an upper limit of 1500 μmol mol^−1^. For each CO_2_ concentration, photosynthetic measurement was completed in 3 min. Using the *A*/*C*_i_ curves, the maximum rates of RuBP regeneration (*J*_max_) and carboxylation (*V*_cmax_) were calculated (Long & Bernacchi 2003).

The quantum yield of PSII photochemistry was calculated as Φ_PSII_ = (*F*_*m*_*’* – *F*_*s*_)/*F*_*m*_*’* (Genty, Briantais & Baker 1989), where *F*_*m*_*’* and *F*_*s*_ represent the maximum and steady-state fluorescence after light adaptation, respectively (Baker 2004). The total electron transport rate through PSII (*J*_PSII_) was calculated as follows (Krall & Edwards 1992):

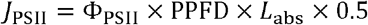

where PPFD is the photosynthetic photon flux density and leaf absorbance (*L*_abs_) is assumed to be 0.84. We applied the constant of 0.5 based on the assumption that photons were equally distributed between PSI and PSII.

### Estimation of mesophyll conductance and chloroplast CO_2_ concentration

Mesophyll conductance was calculated according to the following equation (Harley, Loreto, Di Marco & Sharkey 1992):

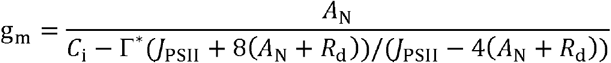

where *A*_N_ represents the net rate of CO_2_ assimilation; *C*_i_ is the intercellular CO_2_ concentration; Γ ^*^ is the CO_2_ compensation point in the absence of daytime respiration (Yamori, Noguchi, Hikosaka & Terashima 2010b; von Caemmerer & Evans 2015), and we used a typical value of 40 μmol mol^-1^ in our current study (Xiong, Douthe & Flexas 2018). Respiration rate in the dark (*R*_d_) was considered to be half of the dark-adapted mitochondrial respiration rate as measured after 10 min of dark adaptation (Carriquí *et al*. 2015).

Based on the estimated *g*_m_, we then calculated the chloroplast CO_2_ concentration (*C*_c_) according to the following equation (Long & Bernacchi 2003; Warren & Dreyer 2006):

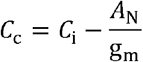

To identify the rate-limiting step of CO_2_ assimilation, we subsequently estimated *C*_trans_ (the chloroplast CO_2_ concentration at which the limitation to *A*_N_ transitioned from RuBP carboxylation to RuBP regeneration) (Yamori, Evans & Von Caemmerer 2010a; Yamori, Nagai & Makino 2011):

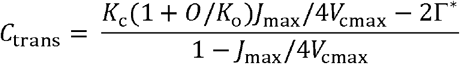

where *K*_c_ (μmol mol^-1^) and *K*_o_ (mmol mol^-1^) are assumed to be 407 μmol mol^-1^ and 277 mmol mol^-1^ at 25ºC, respectively (Long and Bernacchi 2003); *O* was assumed to be 210 mmol mol^-1^ (Farquhar et al. 1980). The rate-limiting step for CO_2_ assimilation was analyzed by comparing the values of *C*_c_ and *C*_trans_. *A*_N_ tends to be limited by RuBP carboxylation when *C*_c_ is lower than *C*_trans_ and tends to be limited by RuBP regeneration when *C*_c_ is higher than *C*_trans_.

### Quantitative limitation analysis of *A*_N_

Relative photosynthetic limitations were assessed as follows (Grassi & Magnani 2005):

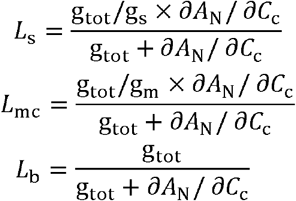

where *L*_s_, *L*_mc_, and *L*_b_ represent the relative limitations of stomatal conductance, mesophyll conductance, and biochemical capacity, respectively, in setting the observed value of *A*_N_. *g*_tot_ is the total conductance of CO_2_ between the leaf surface and sites of RuBP carboxylation (calculated as 1/*g*_tot_ = 1/*g*_s_ + 1/*g*_m_).

### P700 and chlorophyll fluorescence measurements

PSI and PSII parameters were measured at 25ºC under atmospheric CO_2_ condition using a Dual-PAM 100 measuring system (Heinz Walz, Effeltrich, Germany). Light from a 635-nm light-emitting diode array equipped in Dual-PAM 100 was used as actinic light for illumination. After dark adaptation for at least 15 min, a saturating pulse (20,000 μmol photons m^−2^ s^−1^, 300 ms) was used to measure the maximum fluorescence intensity (*F*_*m*_) and the maximum photo-oxidizable P700 (*P*_*m*_). Subsequently, leaves were exposed to 1455 μmol photons m^−2^ s^−1^ for 5 min to activate photosynthetic electron sinks and then illuminated at fluctuating light alternating between low light (59 μmol photons m^−2^ s^−1^, 2 min) and high light (1455 μmol photons m^−2^ s^−1^, 1 min). During this fluctuating light treatment, PSI and PSII parameters were recorded simultaneously. After the fluctuating light treatment, *P*_*m*_ was measured after a 15-min dark adaptation.

The chlorophyll fluorescence parameters were calculated as follows: Y(II) = (*F*_*m*_*’* − *F*_*s*_)/ *F*_*m*_*’*; NPQ = (*F*_*m*_ − *F*_*m*_*’*)/ *F*_*m*_*’*; Y(NO) = *F*_*s*_/*F*_*m*_. Y(II) is the quantum yield of PSII photochemistry; NPQ, non-photochemical quenching in PSII; Y(NO), the quantum yield of non-regulatory energy dissipation in PSII. *F*_*m*_ and *F*_*m*_*’* are the maximum fluorescence intensity after dark and light acclimation, respectively. *F*_*s*_ is the light-adapted steady-state fluorescence.

PSI photosynthetic parameters were measured by a Dual PAM-100 based on P700 signal (difference of intensities of 830 and 875 nm pulse-modulated measuring light reaching the photodetector). The P700^+^ signals (*P*) may vary between a minimal (P700 fully reduced) and a maximal level (P700 fully oxidized). The maximum level (*P*_*m*_) was determined with application of a saturating pulse (20,000 μmol photons m^−2^ s^−1^ and 300 ms) after pre-illumination with far-red light, and *P*_*m*_ was used to estimate the PSI activity. *P*_*m*_*’* was determined similarly to *P*_*m*_ but with actinic light instead of far-red light. PSI parameters were calculated as follows: Y(I) = (*P*_*m*_*’* − *P*)/*P*_*m*_; Y(ND) = *P*/*P*_*m*_; Y(NA) = (*P*_*m*_ − *P*_*m*_*’*)/*P*_*m*_. Y(I) is the quantum yield of PSI photochemistry; Y(ND), the quantum yield of PSI non-photochemical energy dissipation due to donor side limitation; Y(NA), the quantum yield of PSI non-photochemical energy dissipation due to acceptor side limitation. The photosynthetic electron transport rate was calculated as ETRI (or ETRII) = PPFD × Y(I) [or Y(II)] × 0.84 × 0.5, light absorption is assumed to be 0.84 of incident irradiance, and 0.5 is the fraction of absorbed light reaching PSI or PSII. The relative CEF value was measured by the ratio of Y(I) to Y(II) (Grieco *et al*. 2020).

### Statistical analysis

One-way ANOVA and t-tests were used to determine whether significant differences existed between different treatments (α = 0.05). The software SigmaPlot 10.0 was used for graphing and fitting.

## Results

### Effects of drought stress on gas exchange under FL

After water deficit treatment for one week, the maximum quantum yield of PSII after dark adaptation was maintained at approximately 0.84 (data not shown), indicating that PSII activity was not significantly photoinhibited under drought stress. At low light intensity of 100 μmol photons m^−2^ s^−1^, the steady-state net rate of CO_2_ assimilation (*A*_N_) was slightly affected by drought stress. However, drought stress significantly affected steady-state *A*_N_, stomatal conductance (*g*_s_), and mesophyll conductance (*g*_m_) under high light. Furthermore, the photosynthetic induction after transition from low to high light was largely affected by drought stress (Fig. 1A). After transition from 100 to 1500 μmol photons m^−2^ s^−1^ for 1 min, *A*_N_ rapidly increased to 17.5 μmol m^−2^ s^−1^ in control (CK) plants but only increased to 4.0 μmol m^−2^ s^−1^ under drought (Fig. 1A). The time required for *A*_N_ to reach 80% of the maximum value was approximately 2 min in CK plants but 10 min under drought stress (Fig. 1B). Drought stress decreased *g*_s_ under low and high light compared to CK levels (Fig. 1C). Furthermore, the *g*_s_ level at low light was greatly reduced under drought stress compared with the maximum *g*_s_ under high light, whereas there was less relative difference in *g*_s_ between low and high light in CK (Fig. 1D). Drought stress also lowered *g*_m_ under high light compared to CK (Fig. 1E). After transition from 100 to 1500 μmol photons m^−2^ s^−1^, the time taken for *g*_m_ to reach the maximum value was approximately 3 min in CK plants but 11 min under drought stress (Fig. 1F). Therefore, drought stress delayed the induction responses of *g*_s_, *g*_m_, and *A*_N_ under FL.

**Figure 1.**
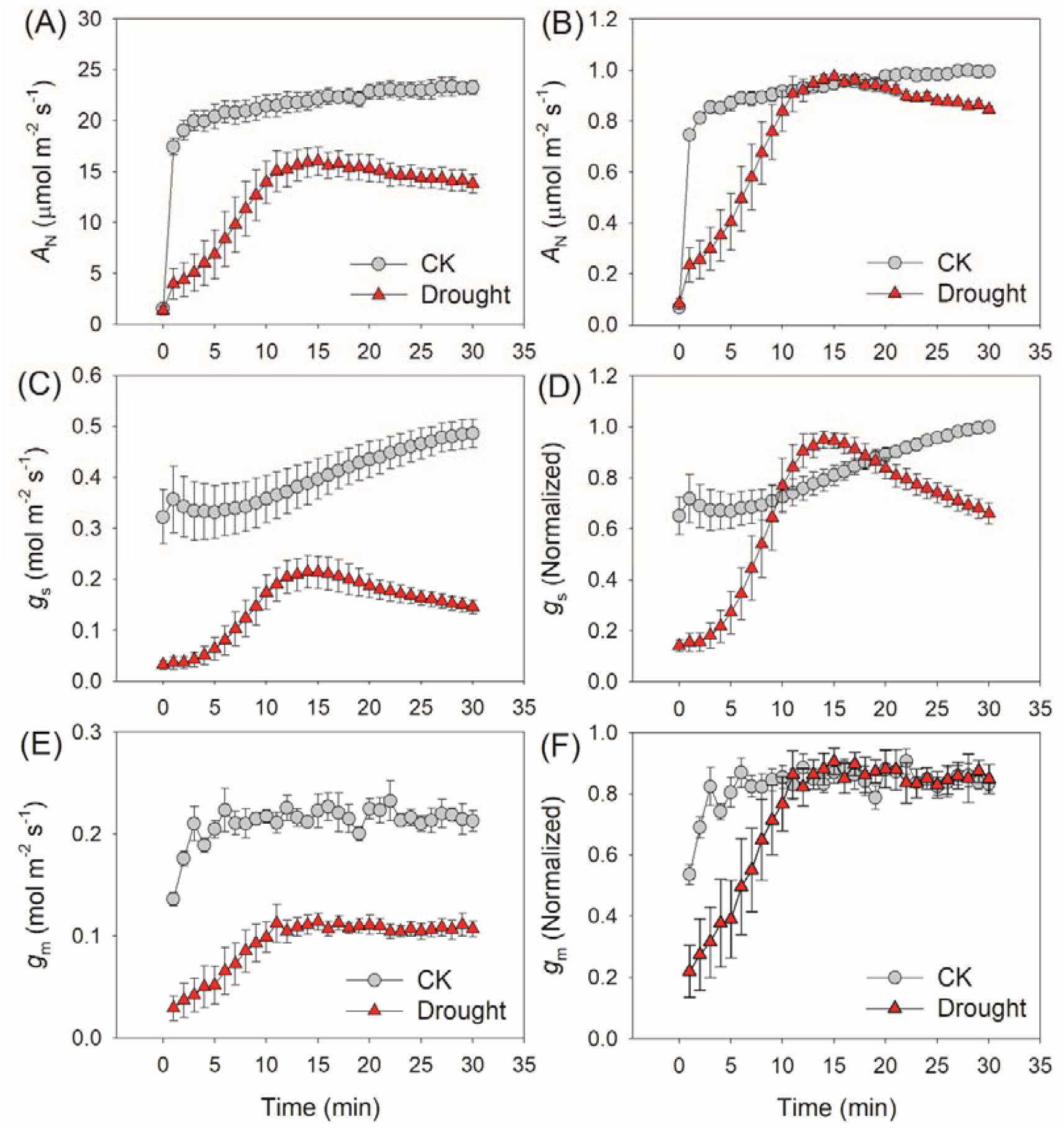
The effects of drought stress on photosynthetic induction after transition from low to high light. Time course of net CO_2_ assimilation (*A*_N_; A and B), stomatal conductance (*g*_s_; C and D) and mesophyll conductance (*g*_m_; E and F) after transition from 100 to 1500 μmol photons m^−2^ s^−1^. Before this measurement, leaves were adapted to low irradiance (100 μmol photons m^−2^ s^−1^) for 5 min. *A*_N_, *g*_s_ and *g*_m_ were normalized against the maximum values after photosynthetic induction for 30 min. Values are means ± SE of five independent experiments (n = 5).

Due to lower levels of CO_2_ diffusional conductance, the intercellular CO_2_ concentration (*C*_i_) and chloroplast CO_2_ concentration (*C*_c_) under high light largely decreased under drought stress compared to CK (Fig. 2A and B). The values of *C*_c_ during photosynthetic induction were always higher than *C*_trans_ (the chloroplast CO_2_ concentration at which the limitation to *A*_N_ transitioned from RuBP carboxylation to RuBP regeneration) in CK plants but were always lower than *C*_trans_ under drought stress, indicating that *A*_N_ tended to be limited by RuBP regeneration in CK plants and by RuBP carboxylation under drought stress. Therefore, the rate-limiting step for *A*_N_ during photosynthetic induction was altered by drought stress. The change in *A*_N_ during photosynthetic induction was tightly and positively correlated to *C*_c_ (Fig. 2C), indicating CO_2_ fixation under FL was largely restricted by *C*_c_. Quantitative analysis of photosynthetic limitations revealed stomatal limitation (*L*_s_) and mesophyll conductance limitation (*L*_mc_) were enhanced under drought stress (Fig. 3A and B). Concomitantly, biochemistry limitation (*L*_b_) decreased under drought stress (Fig. 3C). Therefore, the major limitation of photosynthesis under FL also changed under drought stress.

**Figure 2.**
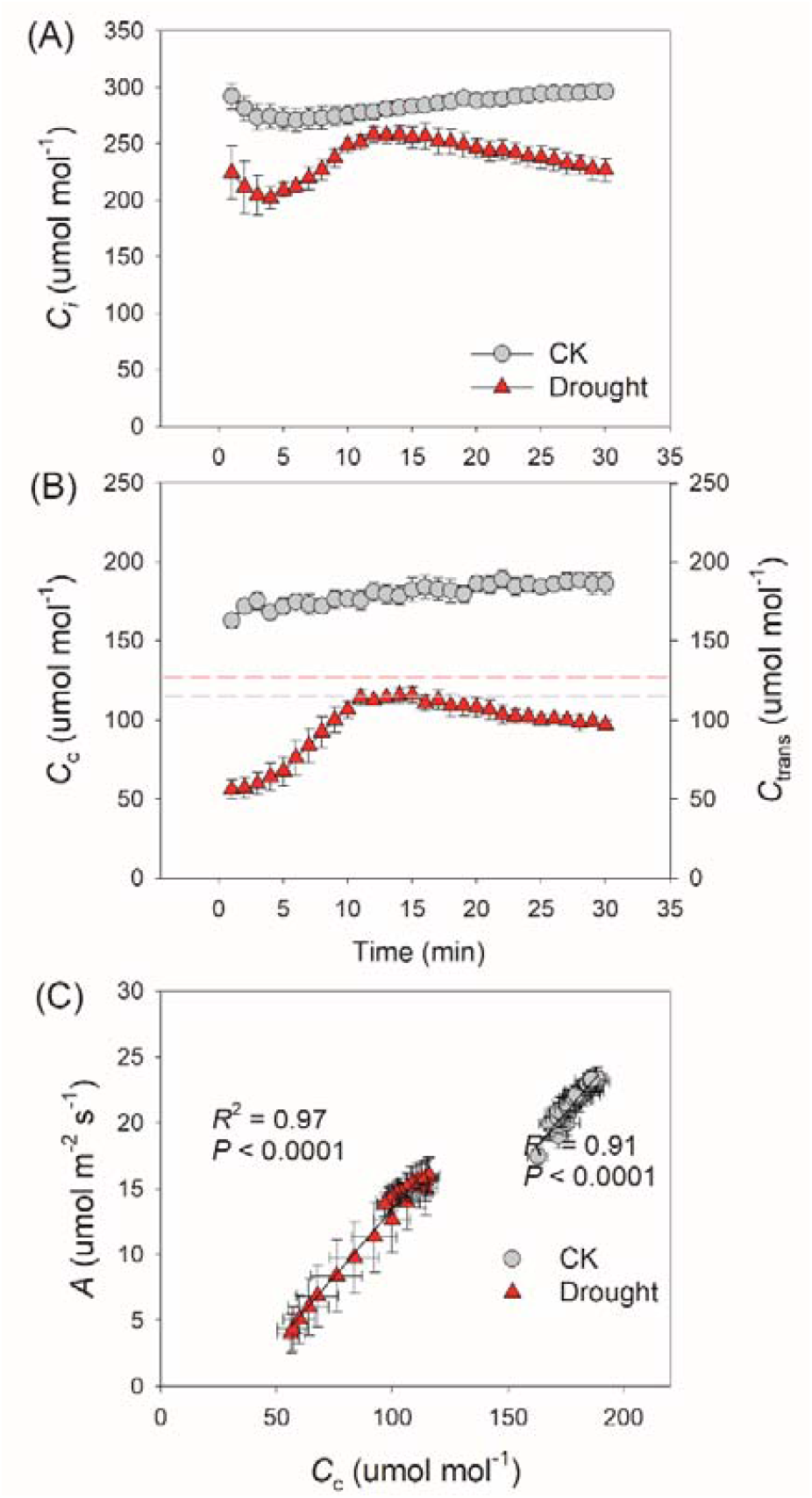
Effects of drought stress on the time course of intercellular CO_2_ concentration (*C*_i_; A) and chloroplast CO_2_ concentration (*C*_c_; B), and the relationship between *C*_c_ and *A*_N_ after transition from low to high light. Red and grey dotted lines represent the values of *C*_trans_ (the chloroplast CO_2_ concentration at which the limitation to *A*_N_ transitioned from RuBP carboxylation to RuBP regeneration) in CK and drought stressed plants, respectively. The experimental design was the same as described in Figure 1. Values are means ± SE of five independent experiments (n = 5).

**Figure 3.**
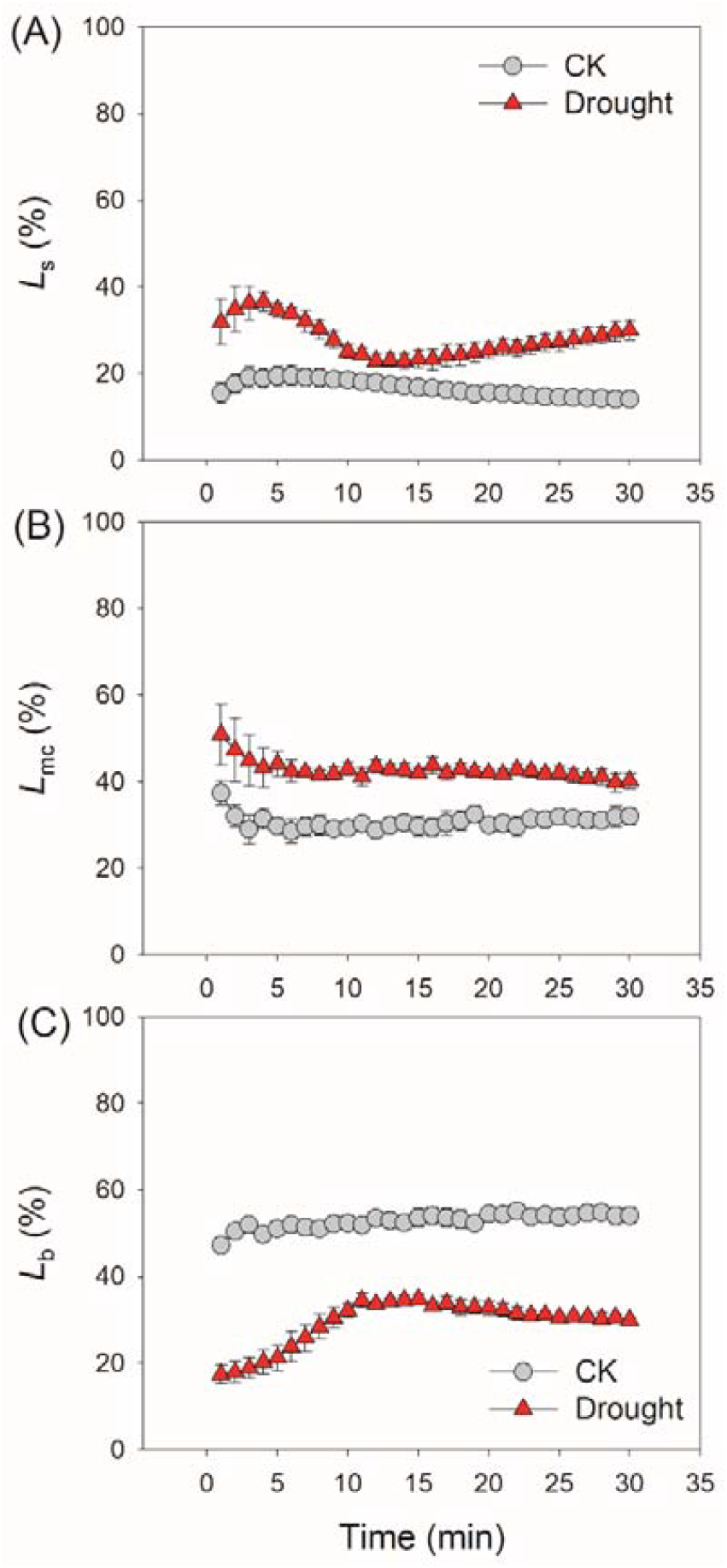
Effects of drought stress on the quantitative analysis of stomatal limitation (*L*_s_), mesophyll conductance limitation (*L*_mc_) and biochemical limitation (*L*_b_) after transition from low to high light. The experimental design was the same as described in Figure 1. Values are means ± SE of five independent experiments (n = 5).

### Effects of drought stress on light reactions under FL

During photosynthetic induction, the quantum yield of PSI photochemistry (Y(I)) under high-light phases decreased under drought stress (Fig. 4A), whereas the PSI donor side limitation (Y(ND)) decreased under drought stress (Fig. 4B), leading to the higher PSI acceptor side limitation (Y(NA)) in plants exposed to drought stress (Fig. 4C). Therefore, drought stress induced stronger PSI over-reduction under FL. Furthermore, under sufficient water supply, the PSI over-reduction under FL was mainly observed after transitioning from low to high light for 10 s (Fig. 4C). By comparison, plants also displayed severe PSI over-reduction after this light transition for 30 s under drought stress (Fig. 4C). Therefore, drought stress prolonged the time course of PSI over-reduction under FL.

**Figure 4.**
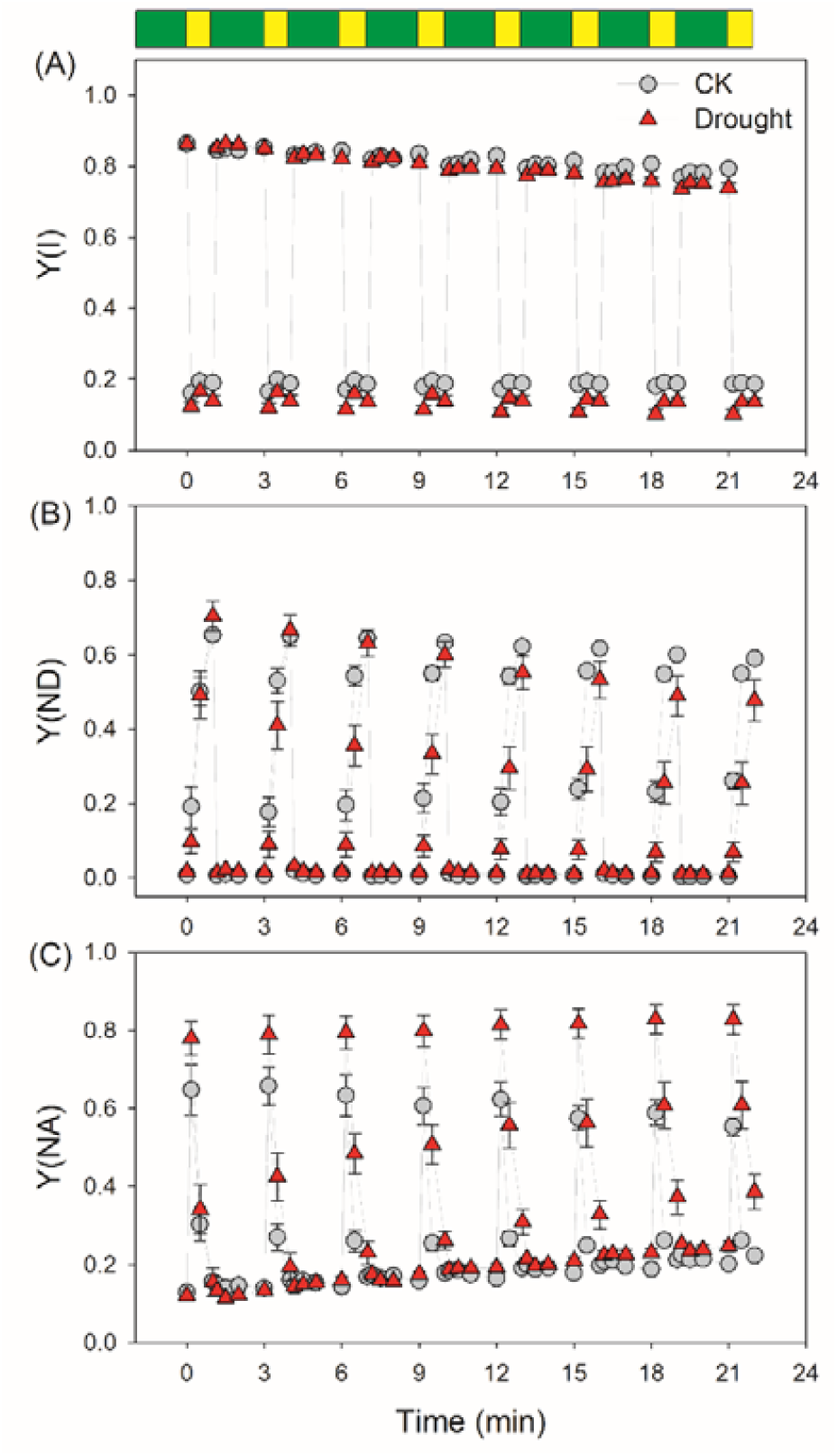
Effects of drought stress on photosystem I parameter under fluctuating light alternating between 59 μmol photons m^−2^ s^−1^ (2 min) and 1455 μmol photons m^−2^ s^−1^ (1 min). Y(I), the quantum yield of PSI photochemistry; Y(ND), the quantum yield of PSI non-photochemical energy dissipation due to donor side limitation; Y(NA), the quantum yield of PSI non-photochemical energy dissipation due to acceptor side limitation. Values are means ± SE of five independent experiments (n = 5).

Similar to Y(I), drought stress decreased the value of Y(II) at high light during photosynthetic induction (Fig. 5A). The induction response of NPQ was not affected by drought stress (Fig. 5B), leading to small changes in the quantum yield of non-regulatory energy dissipation in PSII (Y(NO)) between CK and drought-stressed plants (Fig. 5C). Drought stress had minimal effects on electron transport rates through PSI and PSII (ETRI and ETRII) at low light (Fig. 6A and B). However, the relative values of ETRI and ETRII under high light significantly decreased after drought treatment (Fig. 6A and B), which was consistent with the decreased *A*_N_ under drought stress. The relative CEF value under high light, measured as the ratio of Y(I) to Y(II), was higher in drought-stressed plants compared to CK plants (Fig. 6C). Furthermore, CEF could not be fully activated within the first 10 s under drought stress, suggesting that drought stress induced a delayed activation of CEF under FL. Therefore, the changing model of CEF under FL was altered by drought stress.

**Figure 5.**
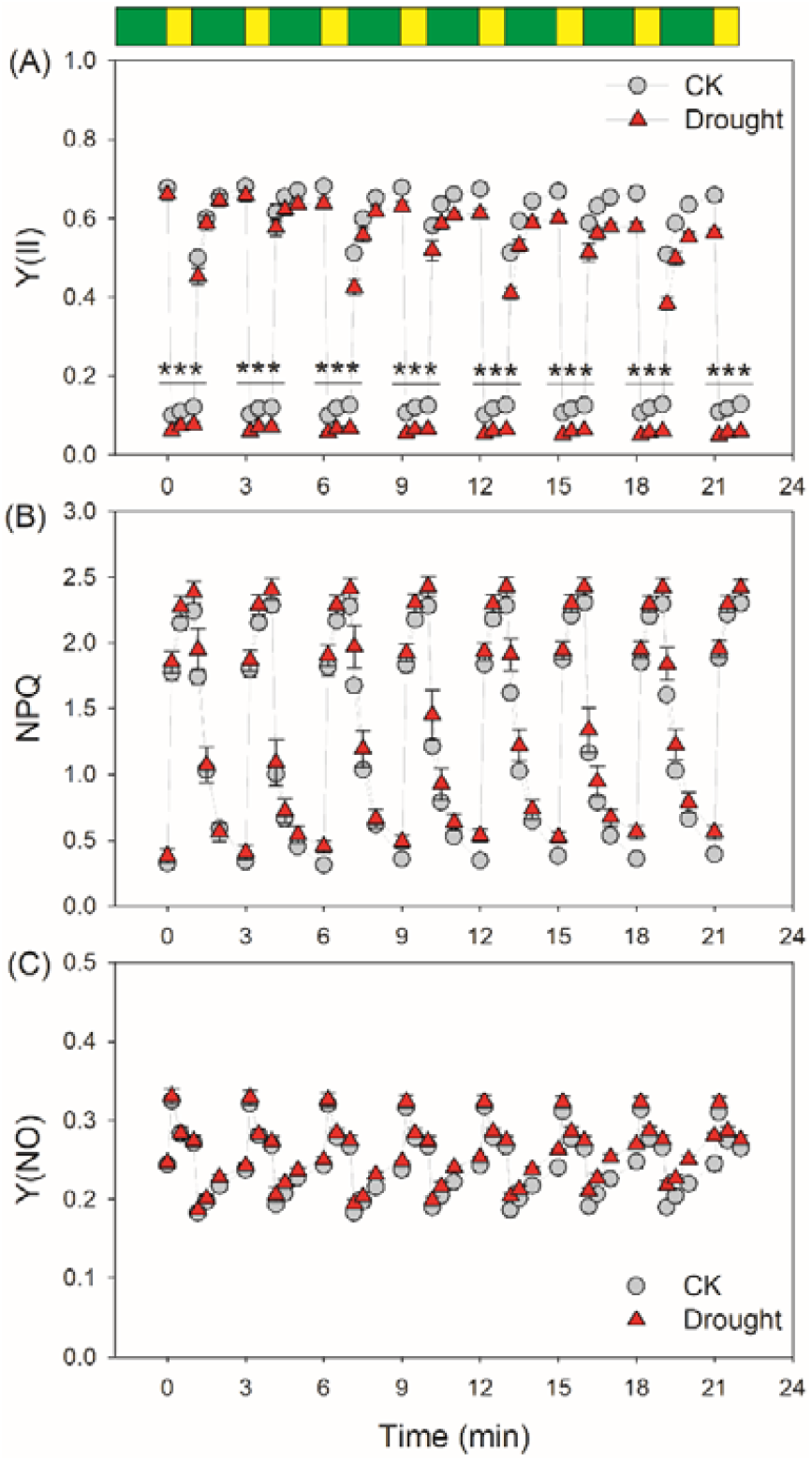
Effects of drought stress on photosystem II parameter under fluctuating light alternating between 59 μmol photons m^−2^ s^−1^ (2 min) and 1455 μmol photons m^−2^ s^−1^ (1 min). Y(II), the quantum yield of PSII photochemistry; NPQ, non-photochemical quenching in PSII; Y(NO), the quantum yield of non-regulatory energy dissipation in PSII. Values are means ± SE of five independent experiments (n = 5).

**Figure 6.**
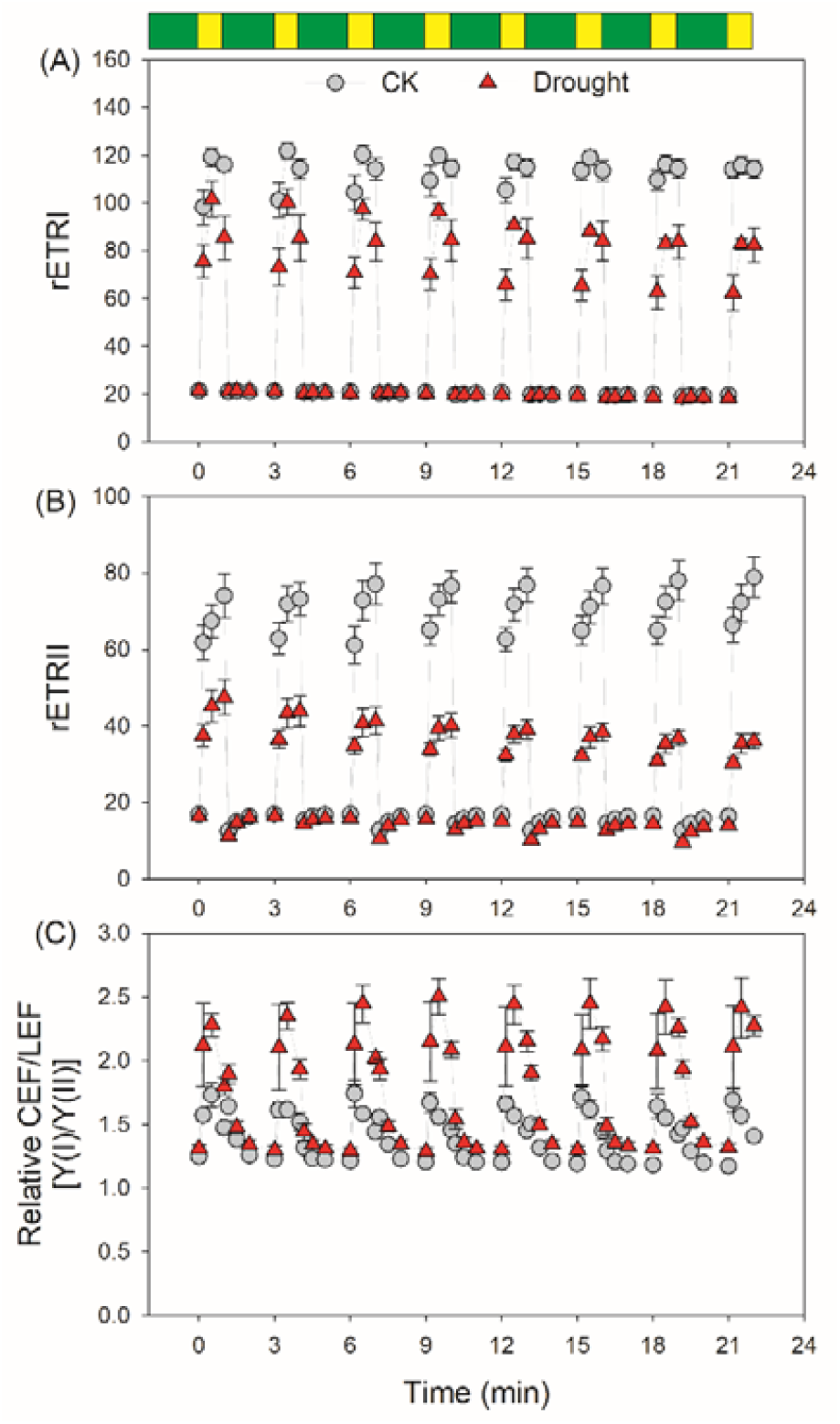
Effects of drought stress on photosynthetic electron transport rate under fluctuating light alternating between 59 μmol photons m^−2^ s^−1^ (2 min) and 1455 μmol photons m^−2^ s^−1^ (1 min). rETRI, the relative electron transport rate through PSI; rETRII, the relative electron transport rate through PSII; Relative CEF/LEF, the relative cyclic and linear electron flow ratio. Values are means ± SE of five independent experiments (n = 5).

After FL treatment, the decrease in *P*_*m*_ was significantly higher under drought stress (Fig. 7A), indicating that drought stress significantly accelerated PSI photoinhibition under FL. The greater FL-induced PSI photoinhibition under drought stress was largely caused by the stronger PSI over-reduction within the first 30 s after transition from low to high light (Fig. 7B). Furthermore, a tight inverse relationship was found between the extent of FL-induced PSI photoinhibition and the maximum CO_2_ assimilation rate (Fig. 7C), suggesting that the restriction of CO_2_ fixation under drought stress accelerated FL-induced PSI photoinhibition. After the FL treatment, ETRII under high light significantly decreased in drought-stressed plants (Fig. 8). Concomitantly, Y(ND) significantly decreased and Y(NA) significantly increased (Fig. 8). These results indicated that the greater FL-induced PSI photoinhibition under drought stress significantly affected linear electron flow and PSI photoprotection under high light.

**Figure 7.**
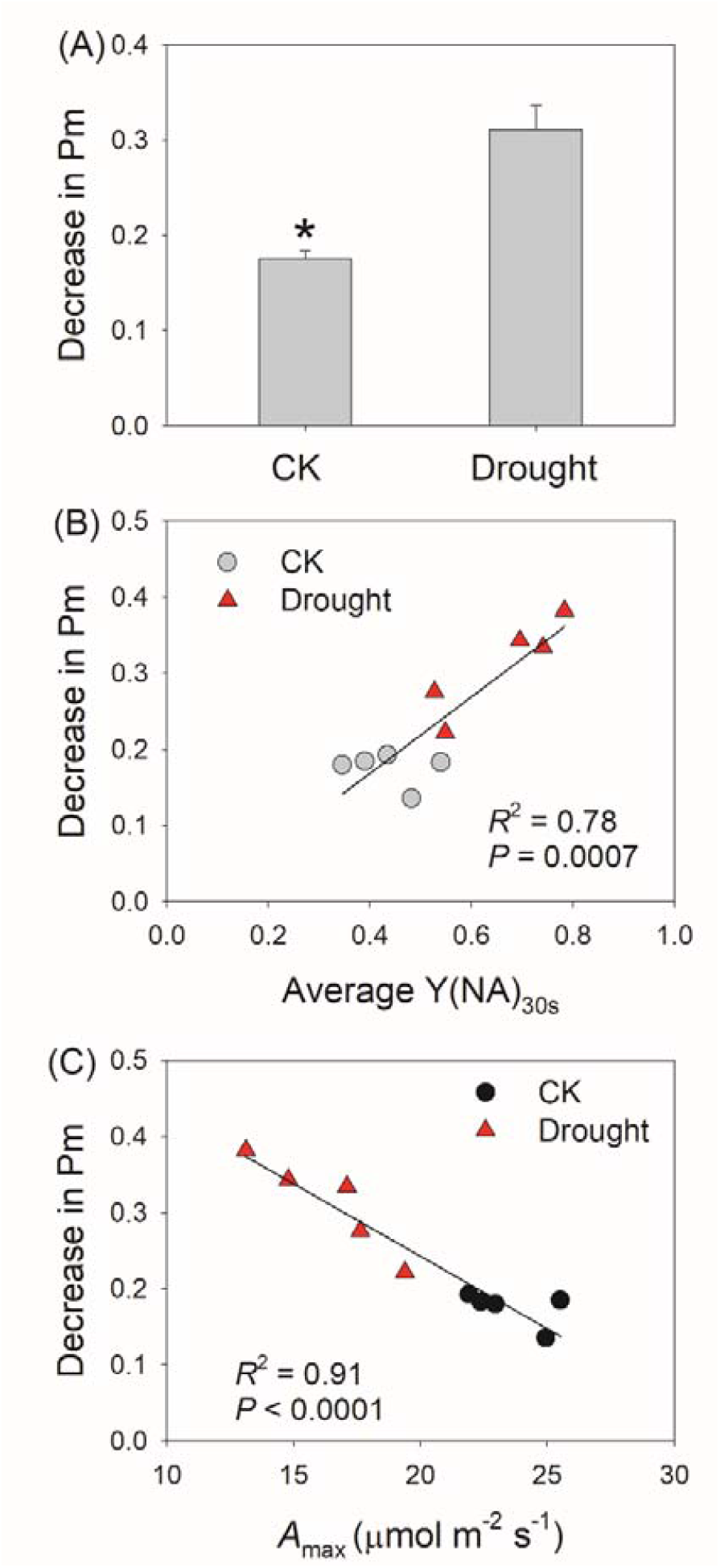
Effects of drought stress on PSI photoinhibition after fluctuating light treatment. (A) The extent of PSI photoinhibition was measured by the decrease in Pm; (B) The relationship between PSI over-reduction within the first 30 s after transition from low to high light [average Y(NA)_30s_] and PSI photoinhibition; (C) The relationship between the maximum CO_2_ assimilation rate during photosynthetic induction (*A*_max_) and PSI photoinhibition. Values are means ± SE of five independent experiments (n = 5). Asterisk indicates a significant different between the CK-plants and drought-stressed plants.

**Figure 8.**
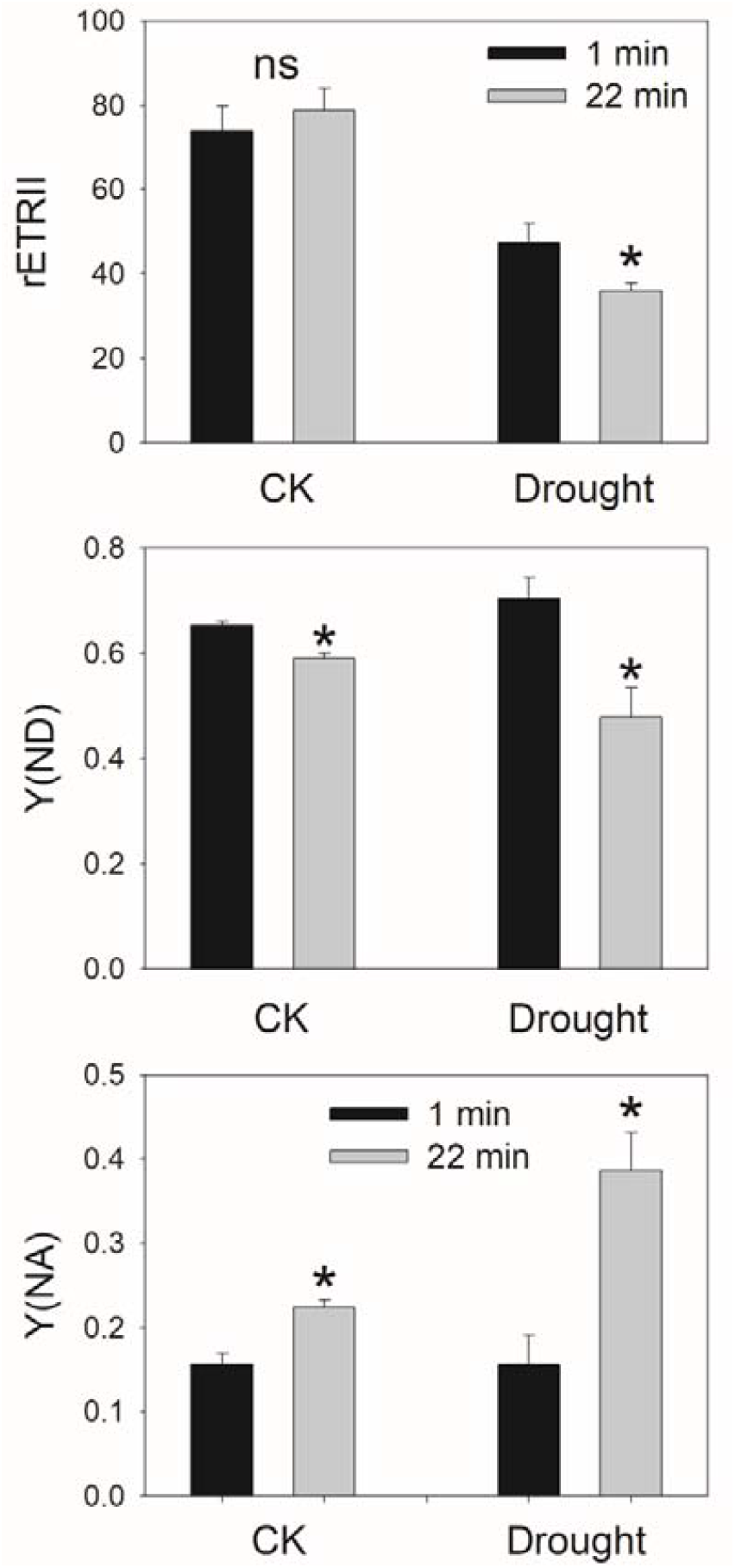
Effects of drought stress on values of rETR, Y(ND) and Y(NA) 1455 μmol photons m^−2^ s^−1^ after fluctuating light treatment. Values are means ± SE of five independent experiments (n = 5). Asterisk indicates a significant different between the CK-plants and drought-stressed plants.

## Discussion

In nature, FL usually occurs concomitantly with drought stress. However, the effects of drought stress on photosynthetic performances under FL are little known. We here for the first time examined the effects of drought stress on photosynthetic induction and PSI photoinhibition under FL in tomato.

### Drought stress delayed photosynthetic induction under FL

The response of *A*_N_ to a rapid change of light intensity plays an important role in determining plant biomass under FL (Vialet-Chabrand, Matthews, Simkin, Raines & Lawson 2017; Slattery *et al*. 2018; Kimura *et al*. 2020; Yamori *et al*. 2020). Drought stress usually decreases steady-state *g*_s_, *g*_m_, and *A*_N_ under constant high light (Warren *et al*. 2004; Huang *et al*. 2013; Zivcak *et al*. 2013), but the effects of drought stress on the induction response of *g*_s_, *g*_m_, and *A*_N_ under FL are not well known. Similar to previous studies, we found the maximum *g*_s_, *g*_m_, and *A*_N_ under constant high light significantly declined in drought-stressed tomato plants (Fig. 1). Moreover, drought stress largely affected the induction responses of *g*_s_ and *g*_m_ after transition from low to high light and consequently delayed the induction response of *A*_N_. *A*_N_ in CK plants reached 80% of the maximum value approximately 2 min after transition from 100 to 1500 μmol photons m^−2^ s^−1^ (Fig. 1). Such photosynthetic induction required 10 min in drought-stressed plants. Therefore, drought stress caused a larger loss of carbon gain upon transition from low to high light in tomato. Similar to observations under drought stress, tomato plants display relatively lower *g*_s_ under salt stress, and the decrease in *g*_s_ under salt stress reduced plant biomass when grown under FL (Zhang, Kaiser, Marcelis, Yang & Li 2020). However, the underlying mechanisms of this response are not well known. Our present study provided a possible explanation for why salt stress reduces plant biomass in tomato plants grown under FL.

Recent studies indicate stomatal opening under FL could be affected by stomatal density and area (Zhang *et al*. 2020; Sakoda *et al*. 2020). In the present study, the drought treatment just lasted for 1 week, and we used mature leaves developed prior to the treatment period for photosynthetic measurements. Therefore, the slower stomatal opening under drought stress was independent of stomatal density and area and was likely caused by other factors such as abscisic acid and gene regulation. Abscisic acid is upregulated under drought stress, which could induce stomatal closure (Ramachandra & Viswanatha 2004; Harb, Krishnan, Ambavaram & Pereira 2010; Kaiser, Morales, Harbinson, Heuvelink & Marcelis 2020). Furthermore, the expression of *slow anion channel-associated 1* (*slac1*), *open stomata 1* (*ost1*), and *proton ATPase translocation control 1* (*PATROL1*) play important roles in stomatal opening under FL (Kimura *et al*. 2020; Yamori *et al*. 2020). Drought stress might influence the expression of these target genes and thus affect the stomatal opening under FL.

Photosynthesis can be limited by *g*_s_, *g*_m_, and biochemical factors, but the relative photosynthetic limitations largely vary among species (Grassi & Magnani 2005; Carriquí *et al*. 2015; Xiong *et al*. 2018). We found that drought stress increased the diffusional limitation of *A*_N_ under FL, and the major limiting factor of photosynthesis was altered by drought stress (Fig. 3). In *Arabidopsis thaliana* and tobacco, CO_2_ diffusional limitation was the major limiting factor of photosynthesis in the initial 10 min after transition from dark to light (Sakoda *et al*. 2021). By comparison, *L*_s_, *L*_mc_ and *L*_b_ changed little after the transition from low to high light in the CK plants (Fig. 3). Therefore, the dynamic changes in relative limitations of *A*_N_ after the transition from low to high light differed from that after the transition from dark to high light. Due to decreased *g*_s_ and *g*_m_ under drought stress, *C*_c_ was lower in high-light phases under FL, and therefore photosynthesis under FL was strongly restricted by *C*_c_ (Fig. 2). Furthermore, after transition from low to high light, the *C*_c_ values in the CK plants were always higher than *C*_trans_ (Fig. 2), indicating that *A*_N_ was mainly limited by RuBP regeneration. By contrast, the *C*_c_ values under FL were always lower than *C*_trans_ under drought stress (Fig. 2), and hence *A*_N_ tended to be mainly limited by RuBP carboxylation. Therefore, the rate-limiting step for *A*_N_ during photosynthetic induction was influenced by drought stress.

### Drought stress accelerated PSI photoinhibition under FL

A sudden increase in light intensity causes a rapid increase in PSII electron flow to PSI, whereas the full activation of CO_2_ fixation requires more time (Lawson & Blatt 2014; Yamori *et al*. 2016). Therefore, the excited states in PSI cannot be immediately consumed by primary metabolism, leading to the accumulation of reducing power in PSI. The resulting PSI over-reduction promotes ROS formation in PSI and causes PSI photoinhibition (Yamori *et al*. 2016; Huang *et al*. 2019a). Under drought stress, lower *g*_s_ restricted CO_2_ fixation and thus decreased the rate of NADPH production. Because the pool of NADPH is relatively small, drought stress will increase the NADPH/NADP^+^ ratio when exposed to a sudden increase in illumination. Under such conditions, electron flow from PSI to NADP^+^ would be limited by the lack of NADP^+^, suppressing the oxidation of PSI under FL (Grieco *et al*. 2020). Therefore, drought stress is hypothesized to accelerate PSI over-reduction under FL. Consistently, we found that PSI over-reduction under FL was aggravated under drought stress (Fig. 4). After a sudden increase in irradiance, tomato plants showed a transient PSI over-reduction in CK plants, indicating that the electrons transported to PSI could not be immediately consumed by downstream sinks. In angiosperms, outflows of electrons from PSI include two pathways: linear electron flow (LEF) (PSI to NADP^+^) and the water-water cycle mediated by the Mehler reaction (Ilík *et al*. 2017; Shikanai & Yamamoto 2017). Recent studies have indicated that the water-water cycle can rapidly consume the excess reducing power in PSI and thus prevents a transient PSI over-reduction under FL (Huang *et al*. 2019b; Sun *et al*. 2020a). However, a transient PSI over-reduction under FL was clearly observed (Fig. 4C), indicating that the water-water cycle is negligible in tomato leaves. Therefore, the transient PSI over-reduction under FL is attributed to the limitation of LEF. Under drought stress, the restriction of CO_2_ fixation caused LEF to be further restricted by the lack of NADP^+^ and thus increased PSI over-reduction upon the transition from low to high light.

Once PSI is over-reduced under high light, the donation of electrons from PSI electron carriers to O_2_ accelerates, aggravating the production of ROS within PSI (Takagi *et al*. 2016). Moreover, ROS produced within thylakoid membranes cannot be immediately scavenged by antioxidant systems (Takagi *et al*. 2016). Therefore, PSI over-reduction under high light easily causes PSI photoinhibition (Yamamoto & Shikanai 2019; Tan *et al*. 2021). Consistently, the stronger PSI over-reduction in FL accelerated PSI photoinhibition under drought stress (Fig. 7). Furthermore, the stronger PSI photoinhibition under drought stress led to decreased rETRII and Y(ND) and increased Y(NA), suggesting that light use efficiency and photoprotection were significantly affected by the PSI photoinhibition (Fig. 8). After photodamage, the recovery of PSI activity is a slow process that requires several days (Zhang & Scheller 2004; Zivcak *et al*. 2015). During the recovery period, the lower CO_2_ assimilation rate impairs starch accumulation and plant growth (Lima-Melo, Gollan, Tikkanen, Silveira & Aro 2019). Therefore, when grown under FL, drought stress would decrease plant biomass partially owing to stronger PSI photoinhibition. In contrast to PSI, the PSII excitation pressure under FL was slightly affected by drought stress (Fig. 5), which was consistent with previous studies reporting that PSII is tolerant to FL (Yamamoto & Shikanai 2019).

To protect PSI against photoinhibition under FL, angiosperms mainly use CEF to fine-tune the PSI redox state through regulation of ΔpH (Armbruster *et al*. 2017). Under low light, upon a low CEF activation, a relatively low ΔpH is formed to facilitate electron flow through Cyt b6/f and thus to maximize light use efficiency (Tikkanen & Aro 2014). Under high light, CEF is highly activated to generate a high ΔpH, which slows down the oxidation of plastoquinone and thus down-regulates the rate of electron transport toward PSI (Shikanai & Yamamoto 2017). When transitioning from low to high light, the full acidification of the thylakoid lumen requires dozens of seconds (Huang *et al*. 2019b; Yang, Zhang & Huang 2019b). Therefore, plants cannot generate sufficient ΔpH within the first seconds after an abrupt increase in irradiance. To avoid uncontrolled PSI over-reduction under FL, CEF rapidly activates to aid the formation of ΔpH (Kono *et al*. 2014; Yang *et al*. 2019a). In the present study, we found relative CEF activity in high-light phases of FL increased under drought stress (Fig. 6), suggesting that the relative contribution of CEF to ΔpH formation was enhanced in drought-stressed plants. Such up-regulation of CEF activity partially compensated for the restriction of CO_2_ fixation and thus favored PSI photoprotection under FL.

## Conclusions

We established that drought stress largely affects *g*_s_ and *g*_m_ after the transition from low to high light and thus delays photosynthetic induction under FL. Furthermore, restriction of CO_2_ assimilation under drought stress accelerates PSI over-reduction under FL, which increases the susceptibility of PSI to photoinhibition. Therefore, drought stress strongly affects the photosynthetic dark and light reactions under FL. These findings provide insight into photosynthetic physiology under drought and FL.

## Acknowledgements

The authors acknowledge the financial support by the National Natural Science Foundation of China (No. 31971412, 32171505) and the Project for Innovation Team of Yunnan Province.

